# Overexpression of HLA class I predominantly on alpha cell in at risk individuals for type 1 diabetes

**DOI:** 10.1101/2020.07.13.201079

**Authors:** Mehdi A. Benkahla, Somayeh Sabouri, William B. Kiosses, Sakthi Rajendran, Estefania Quesada-Masachs, Matthias von Herrath

## Abstract

Human leukocyte antigens of class-I (HLA-I) molecules are hyper-expressed in insulin-containing islets (ICI) of type 1 diabetic (T1D) donors. This study investigated the HLA-I expression in autoantibody positive (AAB+) donors and more closely defined its intra-islet and intracellular localization as well as proximity to infiltrating CD8 T cells with high-resolution confocal microscopy. We found HLA-I hyper-expression had already occurred prior to clinical diagnosis of T1D in islets of AAB+ donors. Interestingly, throughout all stages of disease, HLA-I was mostly expressed by alpha cells. Hyper-expression in AAB+ and T1D donors was associated with intra-cellular accumulation in the Golgi. Proximity analysis showed a moderate but significant correlation between HLA-I and infiltrating CD8 T cells only in ICI of T1D donors, but not in AAB+ donors. These observations not only demonstrate a very early, islet-intrinsic immune-independent increase of HLA-I during diabetes pathogenesis, but also point towards a role for alpha cells in T1D.

**Summary:** Human leukocyte antigens of class-I (HLA-I) are hyper-expressed in insulin-containing islets of type 1 diabetic (T1D) donors. This study finds HLA-I hyper-expression occurs prior to clinical diagnosis of T1D in islets of autoantibody-positive donors and mostly expressed by alpha cells.

## 1 Introduction

Type 1 diabetes (T1D) is an autoimmune disease characterized by increased blood glucose levels likely due to the destruction of insulin-producing beta cells resulting in lifelong dependence on exogenous insulin. This immune attack is mediated by CD8 T cells recognizing beta cell peptides presented by human leukocyte antigen of class I (HLA-I) (Katsarou et al., 2017).

Islet proteins are cleaved into peptides in the proteasome. Peptides are then transported from the cytosol into the endoplasmic reticulum (ER) by the transporter associated with antigen processing (TAP). In the ER, peptides are loaded into HLA-I molecules. The newly formed peptide-HLA-I complex is then discharged in vesicles that migrate to the cell surface through the Golgi apparatus. At the surface, these peptide-HLA-I complexes may be recognized by CD8 T cells (Leone et al., 2013).

T1D is associated with the presence of autoantibodies prior to disease onset. The expansion of autoreactive B cells results in autoantibodies targeting insulin (IAA), glutamate decarboxylase 2 (GAD65), insulinoma-associated protein 2 (IA-2) and zinc transporter 8 (ZnT8) (Krischer et al., 2015). Ziegler et al. showed that children with multiple autoantibodies have greater risk of developing the disease than children with single autoantibody (Ziegler et al., 2013).

Previous studies (Bottazzo et al., 1985, Foulis et al., 1987, Pujol-Borrell et al., 1986, Skog et al., 2015 and Richardson et al., 2016) have shown that HLA-I is hyper-expressed in the insulin-containing islets (ICI) of T1D donors. However, it is not known whether HLA-I hyper-expression is the cause or consequence of T1D pathogenesis. To the best of our knowledge, quantification of HLA-I hyper-expression has not been reported in the islets of AAB+ donors. We therefore characterized the expression and intracellular localization of HLA-I in single and double AAB+ donors as well as in non-diabetic and T1D donors.

Our new findings here show that HLA-I is hyper-expressed intracellularly and accumulates significantly in the Golgi compartments in the AAB+ and T1D donors. Also, HLA-I is mostly expressed on alpha cells, irrespective of disease status. Moreover, a moderate positive correlation between HLA-I expression and the number of CD8 T cells was only observed in the ICIs of T1D donors, but not prior to clinical diabetes in AAB+ donors. These results point towards an early, intrinsic defect of beta and alpha cells in T1D that ultimately might lead to immune recognition.

## 2 Methods

### 2-1 Subjects

Human pancreata collected from cadaveric organ donors were obtained through Network for Pancreatic Organ Donors (nPOD). Six µm thick sections from formalin-fixed paraffin embedded (FFPE) tissue samples from non-diabetic (ND, n = 10), autoantibody positive donors (AAB+, n = 10) and donors with T1D (n = 12) were stained. Donor information is summarized in supplementary Table 1.

### 2-2 Immunofluorescence staining

#### Surface staining

To examine HLA-I (A, B, C) and CD8 T cells expression, pancreatic sections were subject to an immunofluorescence staining as previously described (Sabouri et al., 2020). Briefly, after deparaffinization and rehydration in descending ethanol concentrations, sections were exposed to heat-based citrate antigen retrieval for 20 min. Slides were then incubated with monoclonal mouse anti-human-HLA class I ABC (Abcam, ab70328; 1:200) for 1h at room temperature (RT) followed by detection using polyclonal goat anti-mouse IgG-Alexa Fluor 647 (Life Technologies, 1:1000) for 30 min at RT. Polyclonal rabbit anti-CD8 antibody (ab4055, Abcam, Cambridge MA; 1:400) was incubated overnight at 4 °C, followed by goat anti-rabbit IgG-Alexa Fluor 555 (Life Technologies, 1:1000) for 30 min at RT.

#### Intracellular staining

GM130 was stained using the same protocol with slight modifications. Sections were blocked/permeabilized with 10% goat serum in 0.3% Triton-X-100 PBS for 1 h at RT. Monoclonal rabbit anti-GM130 (EP892Y) antibody,(Abcam ab52649; 1:50) was incubated overnight at 4°C, followed by an Alexa Fluor 647 conjugated goat anti-rabbit IgG antibody (Life Technologies; 1:1000) incubated at RT for 1h in the dark.

### 2-3 Image acquisition and analysis

All 3D images were acquired with a Zeiss laser scanning confocal microscope (LSCM)780 using a 63x (1.4na) objective. All Image stacks (on average 12 slices) were acquired with Nyquist resolution parameters using a 0.3µm step size and optimal frame size of 2644×2644. All 8-bit images were acquired using the full dynamic intensity range (0-256) that was determined with the population of cells having the brightest signals. Samples stained for secondary antibodies and isotypic controls alone were used to define thresholds of real signal above background and non-specific signal. Based on these positive and negative controls, we developed an acquisition strategy that used the same laser power and detector signal amplification (digital gain) settings across all donors. Images were further processed using either Zen Pro (Zeiss) for colocalization or Image Pro Premier (Media Cybernetics) for CD8 T cells proximity analysis. Briefly, maximum intensity projections (MIPs) were generated in Zen and then processed using the Zen colocalization module. Here, regions of interest (ROIs) were drawn around each individual islet based on insulin and glucagon staining. With the inputs of the previously defined minimum thresholds of two fluorescent channels at a time, the software automatically calculated pixel intensity spatial overlap coefficients between them using Mander’s coefficient. Mander’s overlap coefficient is described by MOC = ∑i(Ri x Gi)∑iRi2 x ∑iGi2 where Ri is the intensity of the first (red) fluorophore in an individual pixel, whereas Gi is the corresponding intensity for the second (green) fluorophore in the same pixel (Adler and Parmyrd, 2010). Channel signal comparisons included HLA-I vs insulin, HLA-I vs glucagon and HLA-I vs GM130.

MIPs for CD8 T cell distance localization analysis were created in Image J and further processed with Image Pro Premier. In Image Pro Premier, fluorescently labeled CD8 T cells were automatically detected and outlined as ROIs. These original ROIs were then used to generate expanded concentric radial circles of the original outlined signal. Four x 100 pixel expanded concentric circles were created as shown in Fig. 4b (defined as 100, 200, 300 and 400). These outlined radial circles were then overlaid onto the HLA-I signal images to extract the location, total area and signal intensity of HLA-I that resides within each annulus of concentric circles (See Fig. 4a).

### 2-4 Statistical analysis

Statistical analysis was performed using GraphPad Prism version 8 (GraphPad software, San Diego, CA). Kruskal Wallis was used to calculate p values and Dunn’s multiple comparisons test used as post-hoc test and considered significant if p < 0.05. Correlations were evaluated and considered significant if p < 0.05 and a Pearson’s rank correlation coefficient (r) > 0.70.

## 3 Results

### 3-1 HLA class I is hyper-expressed in the islets of autoantibody positive donors

Pancreas sections from 32 organ donors and a total of 802 islets were examined for the presence of HLA-I by indirect immunofluorescence. Consistent with previous studies, HLA-I was hyper-expressed in the islets of T1D donors and specifically in the insulin-containing islets (ICI) (Fig. 1a). Interestingly, HLA-I was already hyper-expressed in the islets of AAB+ donors (Fig. 1a). ICIs from T1D donors (20.05 ± 12.19 %) showed the highest proportion of HLA-I positive area followed by AAB+ donors (12.63 ± 9.34 %), IDIs from T1D donors (12.29 ± 6.29 %) and ND donors (9.69 ± 5.55 %) (Fig. 1b).

**Figure 1:**
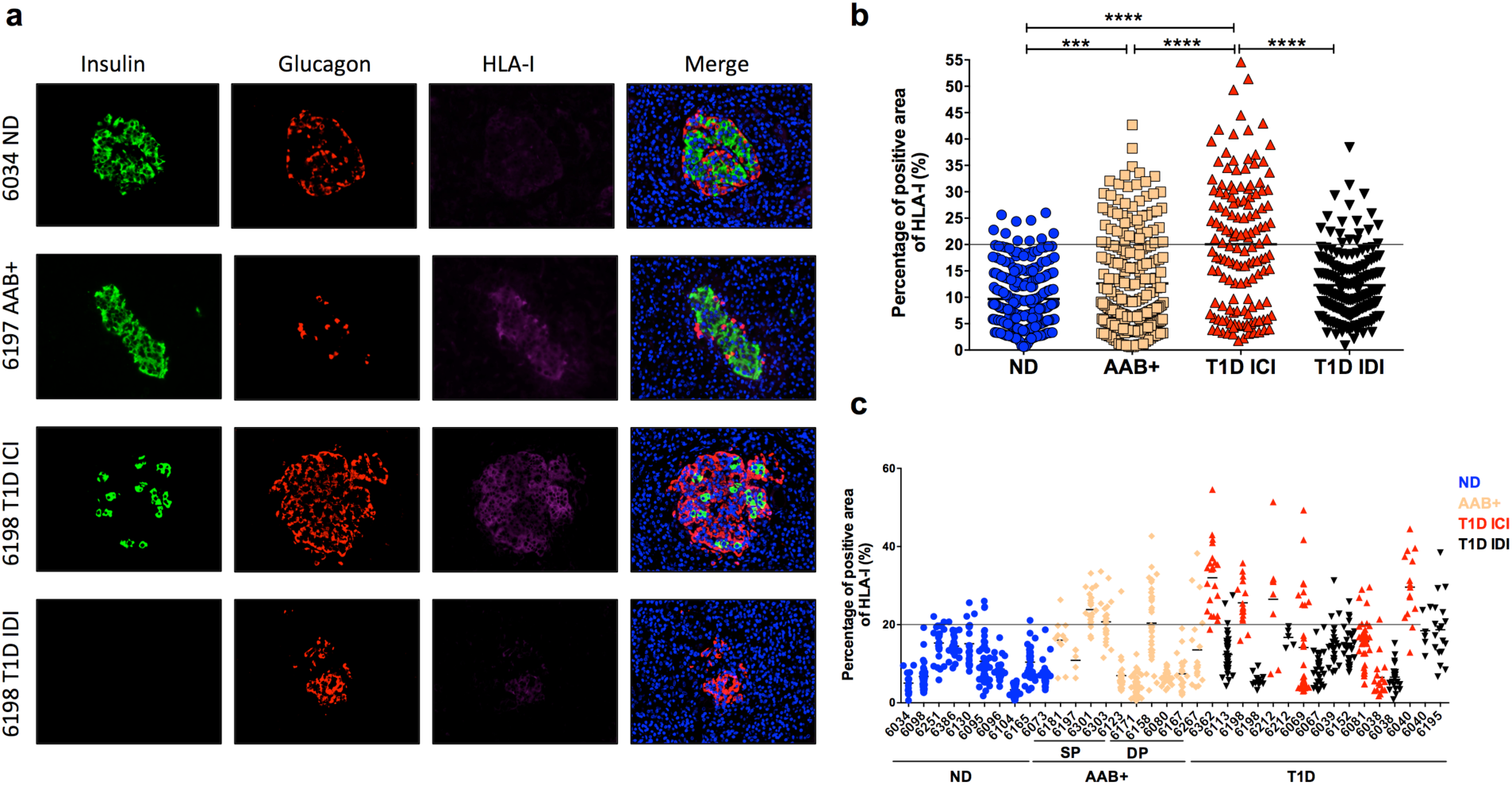
HLA class I is hyper-expressed in the islets of T1D donors as well as in AAB+ donors. FFPE sections of human pancreata were stained with anti-HLA-I (magenta), anti-insulin (green) and anti-glucagon (red) and Hoechst (blue). (a) Representative images of an islet from a non-diabetic (ND) donor (nPOD 6034), an autoantibody positive (AAB+) donor (nPOD 6197), an insulin-containing islet (ICI) of a T1D donor (nPOD 6198) and an insulin-deficient islet (IDI) of a T1D donor (nPOD 6198) are shown. Images were acquired using a confocal microscope LSM 780 with a 63x 1.4na objective. HLA-I was quantified using Zen software. (b) The percentage of total islet area positive for HLA-I was measured in at least 20 islets per section from non-diabetic donors (n=10), single autoantibody positive (n=5), double autoantibody positive (n=5) and type 1 diabetic donors (n=12). (c) Percentage of total islet area positive for HLA-I presented by case. Each data point represents an islet. An arbitrary threshold of 20% was set above which HLA-I was considered hyper-expressed. *** p=0.0004 ****p<0.0001.

The positive area for HLA-I in the ICIs from T1D donors ranged from 1.77 to 54.59 % with 70 islets (out of 142 islets) above the 20% threshold. The positive area for HLA-I in the islets of AAB+ donors ranged from 0.6 to 42.66 % with 52 islets (out of 223 islets) above the 20% threshold. While the positive area for HLA-I for ND donors and IDIs from T1D donors ranged from 0.62 to 26.01 % and 0.84 to 38.41 % respectively, only 10 islets and 19 islets were above the threshold for the ND donors and the IDIs from the T1D donors (Fig. 1b).

HLA-I was hyper-expressed in 52 out of 224 islets (23%) in AAB+ donors (Fig. 1b). A heterogenous expression of HLA-I was observed inter and intra-groups as well as within the same pancreas, where some islets were hyper-expressing HLA-I and others exhibiting a normal HLA-I expression (Fig. 1c). HLA-I hyper-expression was observed not only in recent onset of the disease but also in the islets of donors with T1D for 20 years (nPOD cases 6038, 6040 and 6195) (Fig. 1c).

### 3-2 Majority of HLA class I is expressed on alpha cells

We analyzed the distribution of HLA-I in the islets of ND, AAB+ and T1D donors. The same sections previously analyzed for the quantification of HLA-I were analyzed using Mander’s overlap analysis, to quantify the degree of colocalization of HLA-I with insulin and HLA-I with glucagon. Expression of HLA-I was significantly higher in the alpha cells irrespective of disease status (Fig. 2a). In IDIs, alpha cells contributed to the highest level of HLA-I expression (68.56 ± 14.30 %), which is expected due to the lack of beta cells. Alpha cells in ND, AAB+ and ICIs of T1D donors had a level of distribution of HLA-I of 43.89 ± 25.45 %, 43.35 ± 25.44 % and 53.51 ± 23.80%, respectively.

**Figure 2:**
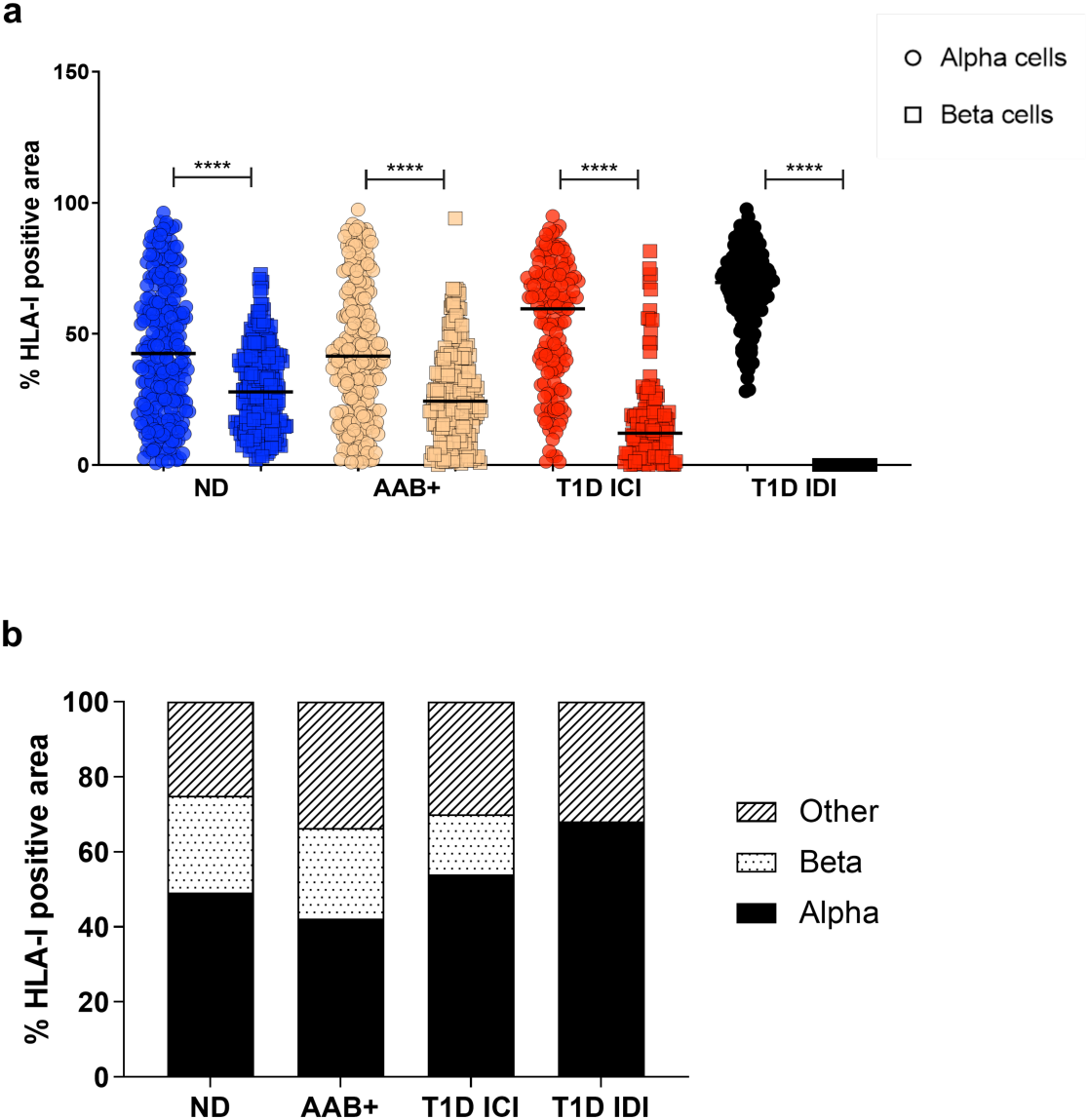
HLA class I is mainly expressed on alpha cells irrespective of disease status. (a) Percentage of colocalization of HLA-I and glucagon (alpha cells-circle) and HLA-I and insulin (beta cells - square) quantified using the Mander’s overlap coefficient using Zen software. Each data point represents an islet. (b) Mean percentage of HLA-I positive area per cell type. ND: non-diabetic; AAB+: autoantibody positive; T1D: type 1 diabetes; ICI: insulin-containing islet; IDI: insulin-deficient islet. **** p<0.0001.

Beta cells in the islets represented the smallest fraction of cells expressing HLA-I in AAB+ and ICIs of T1D donors with a mean percentage of HLA-I expression of 26.79 ± 17.60 % and 16.23 ± 16.51 %, respectively (Fig. 2b).

### 3-3 HLA class I is localized in the Golgi compartment in AAB+ and T1D donors

Next, the intracellular localization of HLA-I was examined. Pancreas section from 1 ND donor, 1 single AAB+ donor, 1 double AAB+ donor and 1 T1D donor were stained for GM130, a peripheral cytoplasmic protein tightly bound to the Golgi membrane, HLA-I, insulin and glucagon. The Mander’s overlap colocalization analysis showed that GM130 and HLA-I were overlapping in the AAB+ and T1D donors with a percentage of colocalization of 7.10 ± 6.56 % in nPOD 6181 AAB+, 5.96 ± 5.29 % in nPOD 6197 AAB+ and 10.95 ± 11.76 % in nPOD 6212 T1D. The non-diabetic donor nPOD 6034 showed the least percentage of colocalization (0.88 ± 0.62 %,Fig. 3).

**Figure 3:**
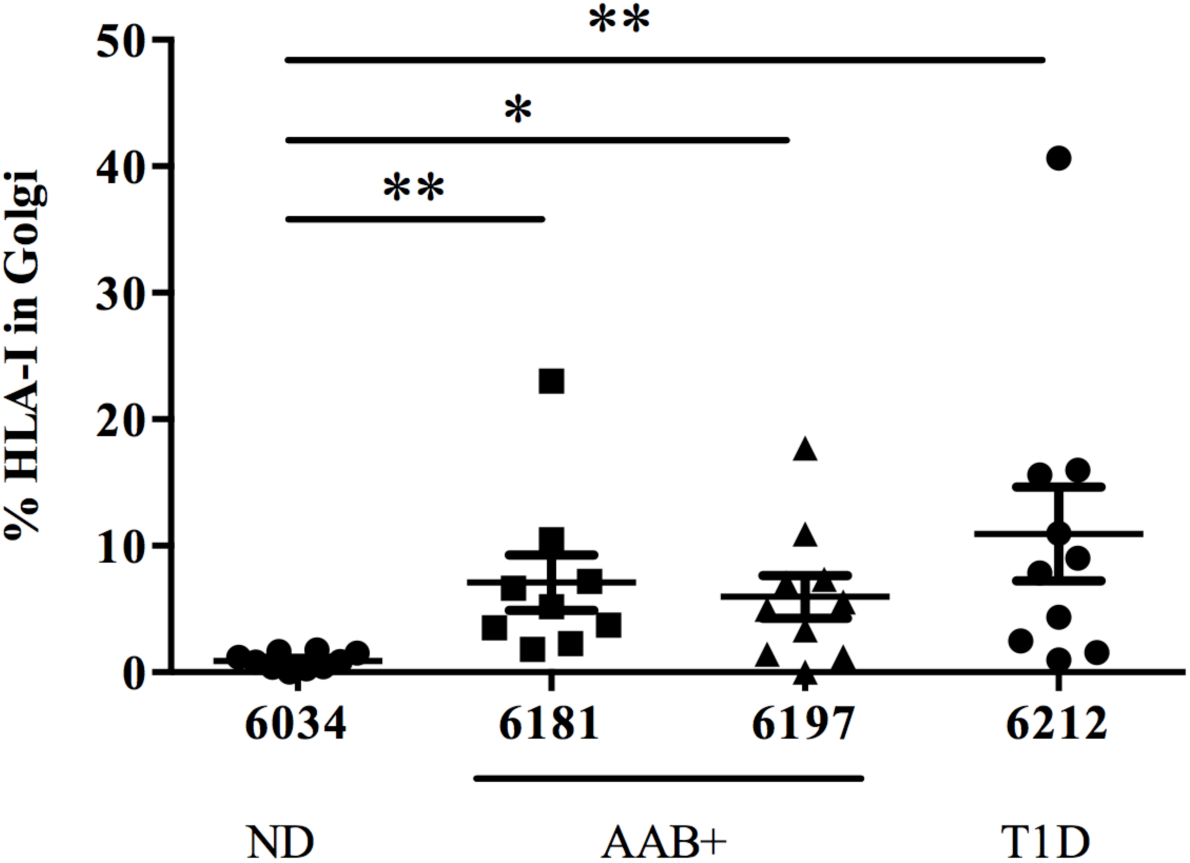
HLA class I colocalizes with Golgi marker GM130 in the islets of AAB+ and T1D donors. Percentage of colocalization of HLA-I and GM130 quantified using Mander’s overlap coefficient in the islets from non-diabetic donor (nPOD 6034), autoantibody positive donors (nPOD 6181 and 6197) and a type 1 diabetic donor (nPOD 6212). Colocalization analysis was performed in Zen software. ND: Non-diabetic; AAB+: autoantibody positive; T1D: type 1 diabetes. *p<0.05 **p<0.01.

### 3-4 HLA class I is hyper-expressed in the proximity of CD8 T cells in T1D but not autoantibody positive donors

CD8 T cells are one of the main perpetrators of the destruction of beta cells through HLA-I (Gomez-Tourino et al 2016). Hence, we examined if the presence of infiltrating CD8 T cells could affect the expression of HLA-I.

Pancreatic sections from 2 ND donors, 1 single AAB+ donor, 1 double AAB+ donor and 2 T1D donors were stained with HLA-I, CD8 and insulin, to examine the expression of HLA-I in the proximity of CD8 T cells. HLA-I expression was invariable between the radial circles surrounding the CD8 T cells irrespective of disease status, but it was higher in the AAB+ and ICIs of T1D donors (Fig. 4a). When examining the whole HLA-I positive area surrounding CD8 T cells (the sum of radial circles), it was significantly higher in the T1D ICIs (Fig 4b,c) compared to the other groups. Data was spread out over a wide range of values with a mean level of HLA-I area of 1597 ± 1469 µm^2^ (Fig. 4c). The levels of HLA-I in the proximity of CD8 T cells ranged from 0 to 4599.7 µm^2^ with a mean of 817.8 ± 991.9 µm^2^ and it was not significantly different between ND donors and IDIs from T1D donors with 631.1 ± 487.7 µm^2^ and 412.5 ± 550.5 µm^2^ respectively (Fig. 4c). Then, we performed a correlation analysis between the number of CD8 T cells and the expression level of HLA-I surrounding those cells. We found a moderate positive correlation between HLA-I and the number of CD8 T cells in the ICIs of T1D with a rho of 0.64 and a p value of 0.016 (Fig. 4d). However, the correlation between HLA-I and the number of CD8 T cells in AAB+ donors was weak with a rho of 0.35 and did not achieve statistical significance with a p value of 0.051.

**Figure 4:**
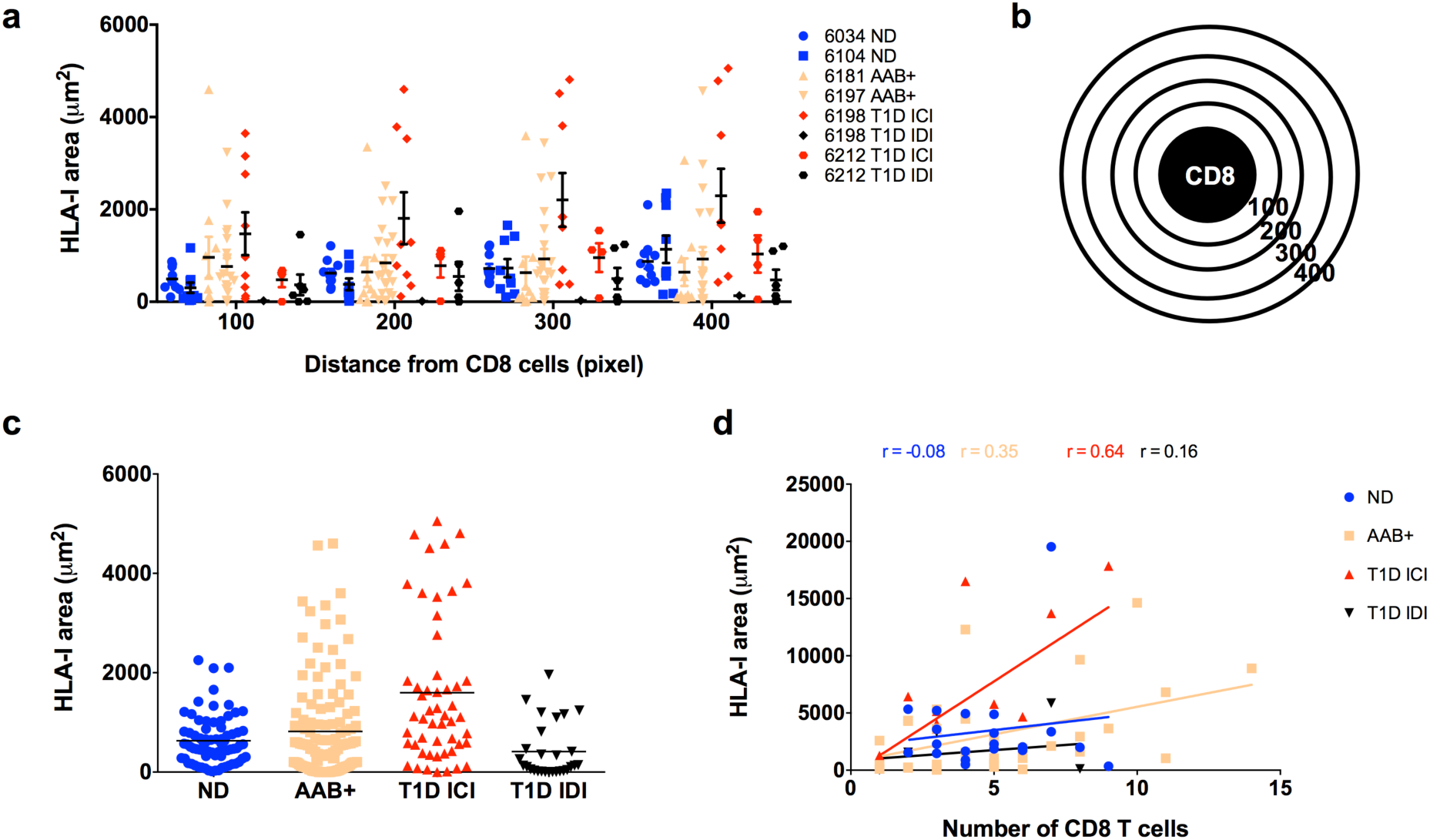
Proximity analysis of HLA class I in the surroundings of CD8 T cells. Pancreatic sections from organ donors were stained for HLA-I, CD8, insulin and glucagon. CD8 T cells were detected in the pancreatic tissue sections and a proximity analysis was performed to detect the HLA-I expression in the surroundings of CD8 T cells. (a) Quantification of HLA-I using ImagePro premier in the radials surrounding the CD8 T cells, 100 being the closest to the CD8 cell and 400 the furthest presented by case. Each data point represents the level of HLA-I in a radial circle. (b) Schematic representation of CD8 T cell and the radial circles from 100 to 400 used for the proximity analysis. (c) Quantification of HLA-I presented as the sum of the radials from 100 to 400 and presented by group. Each data point represents an islet. Each data point represents the level of HLA-I in a radial circle (d) Correlation between the number of CD8 T cells per field of view and the total expression of HLA-I in the surroundings of those CD8 T cells. Each data point represents a field of view. ND: non-diabetic; AAB+: autoantibody positive; T1D: type 1 diabetes; ICI: insulin-containing islet; IDI: insulin-deficient islet.

## Discussion

In agreement with previous studies, HLA-I hyper-expression was observed in insulin-containing islets of T1D donors, confirming that HLA-I hyper-expression is an important pathological feature of type 1 diabetes (Richardson et al., 2016). Skog et al., have reported an intense staining of HLA-I in the islets of AAB+ donors, however, its expression was not characterized nor quantified (Skog et al., 2015).

The current study is different and extends these observations in many aspects: In addition to HLA-I quantification in the islets of ND, AAB+ and T1D donors, its intracellular localization and its possible association with CD8 T cell infiltration have been investigated. Using high-resolution confocal microscopy, we observed a hyper-expression of HLA-I in the insulin-containing islets of T1D donors as well as in the islets of AAB+ donors before the clinical disease onset. In normal conditions, HLA-I molecules are expressed on all cells and are regulated by transcription factors and its expression can be modulated by different stimuli, such as interferon gamma induced in response to viral infections. It has been reported that there was no correlation between HLA-I hyper-expression and enteroviral VP1 (Richardson et al., 2009; Krogvold et al., 2015) and human herpesvirus 6 gB protein in pancreatic islets of T1D donors (Sabouri et al., 2020). Thus, HLA-I upregulation seems to be pointing towards an intrinsic defect with beta and alpha cells preceding and maybe causing T1D.

An interesting new finding in the present study is that the majority of HLA-I is expressed on alpha cells irrespective of disease status. Although expressing higher levels of HLA-I, alpha cells are spared by the immune system compared to beta cells, which are systematically targeted and destroyed. Alpha cells also express other T1D autoantigens such as ZnT8, IA-2 and Chromogranin A, however insulin is exclusively expressed by beta cells, implying that the immune attack of beta cells is related to the presence or presentation of insulin as such (Nakayama, 2011). This might explain why the beta cells are more susceptible to the immune-mediated destruction. Moreover, Brissova et al. have shown that alpha cells from human pancreatic islets obtained from recent-onset T1D donors were functionally impaired and had altered gene expression, which could affect antigen presentation (Brissova et al., 2018).

Furthermore, we have found that HLA-I accumulates in the Golgi apparatus of the islets of AAB+ and T1D donors. This might be explained by HLA-I being hyper-expressed in these donors, resulting in increased intracellular detection. Another possibility could be that ER stress caused by viral infections, environmental stresses or NO (Kataoka and Noguchi, 2013) could lead to accumulation of unfolded or misfolded proteins in the lumen of the ER (Liu and Kaufman, 2003). The latter is further supported by RNA-sequencing analysis performed on alpha cells from T1D donors (Brissova et al., 2018), where an increased expression of genes involved in the unfolded protein response (UPR) was observed. UPR is activated in response to the accumulation of unfolded or misfolded proteins in the ER.

Beta cells are targeted by the CD8 T cell-mediated immune destruction through recognition of autoantigenic peptides associated with HLA-I. HLA-I expression in beta cells is not required for the initiation of insulitis, however it is essential for the destruction of beta cells and incidence of diabetes (Hamilton-Williams et al., 2003). In order to investigate the possible impact of CD8 T cells on HLA-I expression, we performed a proximity analysis of HLA-I in the surroundings of CD8 T cells. HLA-I expression was higher in the surroundings of CD8 T cells in AAB+ and T1D donors, but had normal levels in the ND and IDIs from T1D donors. The limitation of this study is that CD8 T cells are very dynamic, and move around very rapidly as seen in the mouse pancreas by 2-photon imaging (Coppieters et al., 2012). Hence, the proximity analysis may not reflect a true status of CD8 T cell accumulation, although, we have seen a moderate positive correlation between HLA-I expression and the numbers of CD8 T cells in the insulin-containing islets from T1D donors.

In conclusion, our findings have shown that HLA-I is hyper-expressed and accumulates in the Golgi compartment of the islets of AAB+ and T1D donors. Despite being spared by the immune system, alpha cells are the main cell types expressing HLA-I irrespective of disease status. Also, HLA-I is hyper-expressed in the proximity of CD8 T cells in T1D but not AAB+ donors. Taken altogether, our observations points towards an early intrinsic defect in beta and alpha cells that precedes and possibly precipitates immune cell infiltration.

## Author contributions

**MAB:** Data curation, formal analysis, Writing - original draft, Writing - review & editing. **SS:** Data curation, Writing - review & editing. **WBK:** Data curation, Formal analysis, Software, Writing - review & editing. **SR:** Formal analysis, Writing - review & editing. **EQM:** Formal analysis, Writing - review & editing. **MGvH:** Funding acquisition, supervision, formal analysis, writing - review & editing.

## Acknowledgements

This research was performed with the support of nPOD, a collaborative type 1 diabetes research project sponsored by the Juvenile Diabetes Research Foundation International. Organ Procurement Organizations, partnering with nPOD to provide research resources, are listed at www.jdrfnpod.org/our-partners.php. This study was also supported by the National Institutes of Health/National Institute of Allergy and Infectious Diseases grant #U01AI102370-08.

M.A.B. was supported by the Tullie and Rickey SPARK program for innovation in Immunology award from La Jolla Institute for Immunology. S.S. was supported by the NIDDK of the NIH under Award number T32DK007494. EQM was supported by a postdoctoral grant of Fundación Alfonso Martín Escudero.

## References

Adler, J., and I. Parmryd. 2010. Quantifying colocalization by correlation: The Pearson correlation coefficient is superior to the Mander’s overlap coefficient. Cytometry Part A. 77A:733–742. doi: 10.1002/cyto.a.20896.

Bottazzo, G.F., B.M. Dean, J.M. McNally, E.H. MacKay, P.G. Swift, and D.R. Gamble. 1985. In situ characterization of autoimmune phenomena and expression of HLA molecules in the pancreas in diabetic insulitis. N. Engl. J. Med. 313:353–360. doi: 10.1056/NEJM198508083130604.

Brissova, M., R. Haliyur, D. Saunders, S. Shrestha, C. Dai, D.M. Blodgett, R. Bottino, M. Campbell-Thompson, R. Aramandla, G. Poffenberger, J. Lindner, F.C. Pan, M.G. von Herrath, D.L. Greiner, L.D. Shultz, M. Sanyoura, L.H. Philipson, M. Atkinson, D.M. Harlan, S.E. Levy, N. Prasad, R. Stein, and A.C. Powers. 2018. α Cell Function and Gene Expression Are Compromised in Type 1 Diabetes. Cell Rep. 22:2667–2676. doi: 10.1016/j.celrep.2018.02.032.

Coppieters, K., N. Amirian, and M. von Herrath. 2012. Intravital imaging of CTLs killing islet cells in diabetic mice. J Clin Invest. 122:119–131. doi: 10.1172/JCI59285.

Foulis, A.K., M.A. Farquharson, and R. Hardman. 1987. Aberrant expression of class II major histocompatibility complex molecules by B cells and hyperexpression of class I major histocompatibility complex molecules by insulin containing islets in type 1 (insulin-dependent) diabetes mellitus. Diabetologia. 30:333–343. doi: 10.1007/BF00299027.

Gomez-Tourino, I., S. Arif, M. Eichmann, and M. Peakman. 2016. T cells in type 1 diabetes: Instructors, regulators and effectors: A comprehensive review. J. Autoimmun. 66:7–16. doi: 10.1016/j.jaut.2015.08.012.

Hamilton-Williams, E.E., S.E. Palmer, B. Charlton, and R.M. Slattery. 2003. Beta cell MHC class I is a late requirement for diabetes. PNAS. 100:6688–6693. doi: 10.1073/pnas.1131954100.

Kataoka, H.U., and H. Noguchi. 2013. ER Stress and β-Cell Pathogenesis of Type 1 and Type 2 Diabetes and Islet Transplantation. Cell Med. 5:53–57. doi: 10.3727/215517913X666512.

Katsarou, A., S. Gudbjörnsdottir, A. Rawshani, D. Dabelea, E. Bonifacio, B.J. Anderson, L.M. Jacobsen, D.A. Schatz, and Å. Lernmark. 2017. Type 1 diabetes mellitus. Nat Rev Dis Primers. 3:17016. doi: 10.1038/nrdp.2017.16.

Krischer, J.P., K.F. Lynch, D.A. Schatz, J. Ilonen, Å. Lernmark, W.A. Hagopian, M.J. Rewers, J.-X. She, O.G. Simell, J. Toppari, A.-G. Ziegler, B. Akolkar, E. Bonifacio, and TEDDY Study Group. 2015. The 6 year incidence of diabetes-associated autoantibodies in genetically at-risk children: the TEDDY study. Diabetologia. 58:980–987. doi: 10.1007/s00125-015-3514-y.

Krogvold, L., B. Edwin, T. Buanes, G. Frisk, O. Skog, M. Anagandula, O. Korsgren, D. Undlien, M.C. Eike, S.J. Richardson, P. Leete, N.G. Morgan, S. Oikarinen, M. Oikarinen, J.E. Laiho, H. Hyöty, J. Ludvigsson, K.F. Hanssen, and K. Dahl-Jørgensen. 2015. Detection of a Low-Grade Enteroviral Infection in the Islets of Langerhans of Living Patients Newly Diagnosed With Type 1 Diabetes. Diabetes. 64:1682–1687. doi: 10.2337/db14-1370.

Leone, P., E.-C. Shin, F. Perosa, A. Vacca, F. Dammacco, and V. Racanelli. 2013. MHC Class I Antigen Processing and Presenting Machinery: Organization, Function, and Defects in Tumor Cells. J Natl Cancer Inst. 105:1172–1187. doi: 10.1093/jnci/djt184.

Liu, C.Y., and R.J. Kaufman. 2003. The unfolded protein response. Journal of Cell Science. 116:1861–1862. doi: 10.1242/jcs.00408.

Nakayama, M. 2011. Insulin as a key autoantigen in the development of type 1 diabetes. Diabetes Metab Res Rev. 27:773–777. doi: 10.1002/dmrr.1250.

Pujol-Borrell, R., I. Todd, M. Doshi, D. Gray, M. Feldmann, and G.F. Bottazzo. 1986. Differential expression and regulation of MHC products in the endocrine and exocrine cells of the human pancreas. Clin. Exp. Immunol. 65:128–139.

René, C., C. Lozano, and J.-F. Eliaou. 2016. Expression of classical HLA class I molecules: regulation and clinical impacts: Julia Bodmer Award Review 2015. HLA. 87:338–349. doi: 10.1111/tan.12787.

Richardson, S.J., T. Rodriguez-Calvo, I.C. Gerling, C.E. Mathews, J.S. Kaddis, M.A. Russell, M. Zeissler, P. Leete, L. Krogvold, K. Dahl-Jørgensen, M. von Herrath, A. Pugliese, Atkinson, and N.G. Morgan. 2016. Islet cell hyperexpression of HLA class I antigens: a defining feature in type 1 diabetes. Diabetologia. 59:2448–2458. doi: 10.1007/s00125-016-4067-4.

Richardson, S.J., A. Willcox, A.J. Bone, A.K. Foulis, and N.G. Morgan. 2009. The prevalence of enteroviral capsid protein vp1 immunostaining in pancreatic islets in human type 1 diabetes. Diabetologia. 52:1143–1151. doi: 10.1007/s00125-009-1276-0.

Sabouri, S., M.A. Benkahla, W.B. Kiosses, T. Rodriguez-Calvo, J. Zapardiel-Gonzalo, E. Castillo, and M.G. von Herrath. 2020. Human herpesvirus-6 is present at higher levels in the pancreatic tissues of donors with type 1 diabetes. Journal of Autoimmunity. 107:102378. doi: 10.1016/j.jaut.2019.102378.

Skog, O., S. Korsgren, A. Wiberg, A. Danielsson, B. Edwin, T. Buanes, L. Krogvold, O. Korsgren, and K. Dahl-Jørgensen. 2015. Expression of human leukocyte antigen class I in endocrine and exocrine pancreatic tissue at onset of type 1 diabetes. Am. J. Pathol. 185:129–138. doi: 10.1016/j.ajpath.2014.09.004.

Ziegler, A.G., M. Rewers, O. Simell, T. Simell, J. Lempainen, A. Steck, C. Winkler, J. Ilonen, R. Veijola, M. Knip, E. Bonifacio, and G.S. Eisenbarth. 2013. Seroconversion to multiple islet autoantibodies and risk of progression to diabetes in children. JAMA. 309:2473–2479. doi: 10.1001/jama.2013.6285.

