## Supplemental Table 1 for "Overexpression of HLA class I predominantly on alpha cell in at risk individuals for type 1 diabetes"

**Table 1. Demographic and Clinical Information of Pancreatic Organ Donors.**

| **nPOD case number** | **Diagnosis** | **Auto-antibody status** | **Age** | **Sex** | **Race** | **BMI** | **Diabetes duration** | **C-peptide** |
| --- | --- | --- | --- | --- | --- | --- | --- | --- |
| 6073 | Non-diabetic | Negative | 19.2 | Male | Caucasian | 36 | NA | 0.69 |
| 6095 | Non-diabetic | Not tested | 40 | Male | Hispanic/Latino | 35.5 | NA | Not available |
| 6096 | Non-diabetic | Negative | 16 | Female | African Am | 18.8 | NA | 2.97 |
| 6098 | Non-diabetic | Negative | 17.8 | Male | Caucasian | 22.8 | NA | 1.41 |
| 6102 | Non-diabetic | Negative | 45.1 | Female | Caucasian | 35.1 | NA | 0.55 |
| 6104 | Non-diabetic | Negative | 41 | Male | Caucasian | 20.5 | NA | 20.55 |
| 6130 | Non-diabetic | Negative | 5.2 | Male | Caucasian | 18.5 | NA | 4.8 |
| 6165 | Non-diabetic | Negative | 45.8 | Female | Caucasian | 25 | NA | 4.45 |
| 6251 | Non-diabetic | Negative | 33 | Female | Caucasian | 29.5 | NA | 1.92 |
| 6386 | Non-diabetic | Negative | 14 | Male | Caucasian | 23.9 | NA | 1.12 |
| 6080 | AAB+ | GADA and mIAA | 69.2 | Female | Caucasian | 21.3 | NA | 1.84 |
| 6123 | AAB+ | GADA | 23.2 | Female | Caucasian | 17.6 | NA | 2.01 |
| 6158 | AAB+ | GADA | 40.3 | Male | Caucasian | 29.7 | NA | 0.51 |
| 6167 | AAB+ | IA2A and ZnT8 | 37 | Male | Caucasian | 26.3 | NA | 5.43 |
| 6171 | AAB+ | GADA | 4.4 | Female | Caucasian | 14.8 | NA | 8.95 |
| 6181 | AAB+ | GADA | 31.9 | Male | Caucasian | 21.9 | NA | 0.06 |
| 6197 | AAB+ | GADA | 22 | Male | African Am | 28.2 | NA | 17.48 |
| 6267 | AAB+ | GADA and IA2A | 23 | Female | Caucasian | 23.5 | NA | 16.59 |
| 6301 | AAB+ | GADA | 26 | Male | African Am | 32.1 | NA | 3.92 |
| 6303 | AAB+ | GADA | 22 | Male | Caucasian | 31.9 | NA | 3.03 |
| 6038 | T1D | negative | 37.2 | Female | Caucasian | 30.9 | 20 | 0.2 |
| 6039 | T1D | GADA, IA2A, mIAA and ZnT8A | 28.7 | Female | Caucasian | 23.4 | 12 | less than 0.05 |
| 6040 | T1D | mIAA | 50 | Female | Caucasian | 31.6 | 20 | less than 0.05 |
| 6067 | T1D | Negative | 32.6 | Female | Hispanic/Latino | 26.8 | 8 | less than 0.05 |
| 6069 | T1D | Not tested | 22.9 | Male | African Am | 28.8 | 7 | Not available |
| 6081 | T1D | Negative | 31.4 | Male | Hispanic/Latino | 28.0 | 15 | 0.24 |
| 6084 | T1D | mIAA | 14.2 | Male | Caucasian | 26.3 | 4 | less than 0.05 |
| 6113 | T1D | mIAA | 13.1 | Female | Caucasian | 24.8 | 1.6 | less than 0.05 |
| 6152 | T1D | ZnT8A | 29.6 | Female | Caucasian | 30.1 | 12 | less than 0.05 |
| 6195 | T1D | GADA, IA2A, mIAA and ZnT8A | 19.3 | Male | Caucasian | 23.7 | 5 | less than 0.05 |
| 6198 | T1D | GADA, IA2A, mIAA and ZnT8A | 22 | Female | Hispanic/Latino | 23.1 | 3 | less than 0.05 |
| 6212 | T1D | mIAA | 20 | Male | Caucasian | 29.1 | 5 | less than 0.05 |
| 6362 | T1D | GADA | 24.9 | Male | Caucasian | 28.5 | 0 | 0.38 |

AAB+, auto-antibody positive; GADA, GAD autoantibody; IA-2A, Insulinoma-2-associated autoantibody; mIAA, microinsulin autoantibody; ZnT8A, Zinc transporter-8 autoantibodies; NA, not applicable; T1D, type 1 diabetes
